# Deletion of genes linked to the C_1_-fixing gene cluster affects growth, by-products, and proteome of *Clostridium autoethanogenum*

**DOI:** 10.1101/2023.01.30.526161

**Authors:** Ugochi Jennifer Nwaokorie, Kristina Reinmets, Lorena Azevedo de Lima, Pratik Rajendra Pawar, Kurshedaktar Majibullah Shaikh, Audrey Harris, Michael Köpke, Kaspar Valgepea

**Affiliations:** ERA Chair in Gas Fermentation Technologies, Institute of Technology, University of Tartu, 50411 Tartu, Estonia; LanzaTech Inc., 60077 Skokie, USA

**Keywords:** gas fermentation, acetogen, CRISPR/nCas9, genetic engineering, metabolomics, proteomics, chemostat, continuous cultivation, syngas

## Abstract

Gas fermentation has emerged as a sustainable route to produce fuels and chemicals by recycling inexpensive one-carbon (C_1_) feedstocks from gaseous and solid waste using gas-fermenting microbes. Currently, acetogens that utilise the Wood-Ljungdahl pathway to convert carbon oxides (CO and CO_2_) into valuable products are the most advanced biocatalysts for gas fermentation. However, our understanding of the functionalities of the genes involved in the C_1_-fixing gene cluster and its closely-linked genes is incomplete. Here, we investigate the role of two genes with unclear functions – hypothetical protein (*hp;* LABRINI_07945) and CooT nickel binding protein (*nbp;* LABRINI_07950) – directly adjacent and expressed at similar levels to the C_1_-fixing gene cluster in the gas-fermenting model-acetogen *Clostridium autoethanogenum*. Targeted deletion of either the *hp* or *nbp* gene using CRISPR/nCas9, and phenotypic characterisation in heterotrophic and autotrophic batch and autotrophic bioreactor continuous cultures revealed significant growth defects and altered by-product profiles for both Δ*hp* and Δ*nbp* strains. Variable effects of gene deletion on autotrophic batch growth on rich or minimal media suggest that both genes affect the utilisation of complex nutrients. Autotrophic chemostat cultures showed lower acetate and ethanol production rates and higher carbon flux to CO_2_ and biomass for both deletion strains. Additionally, proteome analysis revealed that disruption of either gene affects the expression of proteins of the C_1_-fixing gene cluster and ethanol synthesis pathways. Our work contributes to a better understanding of genotype-phenotype relationships in acetogens and offers engineering targets to improve carbon fixation efficiency in gas fermentation.

## 1. INTRODUCTION

The high concentration of greenhouse gases (GHG) in the atmosphere is leading to potentially irreversible climate change threatening Earth’s sustainability. Fossil fuel combustion for energy, heat, and transportation is widely recognised as the primary source of GHG (mainly CO_2_) from human activities (Sixth Assessment Report | IPCC, 2021). Likewise, increasing population leads to higher volume of generated waste (e.g. plastic waste and municipal solid waste [MSW]) that is negatively impacting ecosystems. Although decarbonising energy generation is important to lowering GHG emissions, we also need to adopt sustainable technologies for the production of fuels and chemicals as these will mostly stay carbon-based for the foreseeable future. The advancement of such technologies will be vital in achieving the United Nations Sustainable Development Goal (SDG) 7 - Affordable and Clean Energy and SDG 13 - Climate Action by the target year 2050.

Gas fermentation has emerged as an attractive route for sustainable production of low-carbon fuels and chemicals from gaseous one-carbon (C1) waste feedstocks (e.g. industrial waste gases [CO_2_, CO, and CH_4_] and syngas (from gasified biomass or MSW [CO, H_2_, and CO_2_]) using gas-fermenting microbes (Fackler et al., 2021; Liew et al., 2016; Pavan et al., 2022; Redl et al., 2017). Anaerobic acetogens are the most advanced biocatalysts for gas fermentation as they can naturally use gas as their sole energy and carbon source (Drake et al., 2006). Acetogens use the most energy-efficient CO_2_ fixation pathway, the Wood-Ljungdahl pathway (WLP), to fix carbon oxides (CO_2_ and/or CO) into many metabolic products via acetyl-CoA. Notably, the model-acetogen *Clostridium autoethanogenum* is being used as a commercial-scale cell factory for ethanol production (Köpke and Simpson, 2020) and has been demonstrated at pilot scale to produce two valuable industrial solvents: acetone and isopropanol (Liew et al., 2022).

The WLP is considered to be the first biochemical pathway on Earth (Russell & Martin, 2004) and it plays a vital role in global carbon cycles by converting ∼20% of CO_2_ on Earth each year to billions of tons of acetate (Drake et al., 2006; Ljungdahl, 2009). The genes encoding the enzymes involved in the WLP belong to a large C1-fixing gene cluster (Figure 1) that has been annotated as CAETHG_1606-1621 (Brown et al., 2014). The authors proposed that the cluster contains 16 genes, including the WLP central enzyme – the bifunctional CO dehydrogenase/acetyl-CoA synthase (CODH/ACS; CAETHG_1620-21/1608) complex – that catalyses CO oxidation/CO_2_ reduction and acetyl-CoA synthesis (Ragsdale and Pierce, 2008) and is essential for autotrophic growth (F. Liew et al., 2016). The cluster also encodes the pathway’s most abundant but slowest enzyme in *C. autoethanogenum* (Valgepea et al., 2022): the formyl-THF ligase (Fhs; CAETHG_1618). Though the biochemistry of the WLP is well-described (Can et al., 2014; Ragsdale, 2008; Ragsdale and Pierce, 2008), our understanding of the functionalities of genes within the proposed C1-fixing gene cluster is incomplete. For instance, functions of genes CAETHG_1607 and 1613 are unclear in *C. autoethanogenum*. In fact, the C1-fixing gene cluster might be larger as several adjacent genes with unknown functions show similarly high gene and protein expression levels (Valgepea et al., 2017, 2022). Two of such genes, found right downstream of the currently annotated cluster, are hypothetical protein (*hp*; CAETHG_1604) and CooT family nickel binding protein (*nbp*; CAETHG_1605). Both *nbp* and *hp* genes and proteins are expressed highly in *C. autoethanogenum* and their homologs accompany the WLP genes of other commonly studied acetogens like *C. ljungdahlii, C. drakei, C. carboxidivorans, Eubacterium limosum*, as well as thermophilic acetogens *Acetobacterium woodii* and *Thermoanaerobacter kivui*. Despite the clear genomic conservation of the *hp* and *nbp* genes among acetogens, their function has not yet been described.

**Figure 1.**
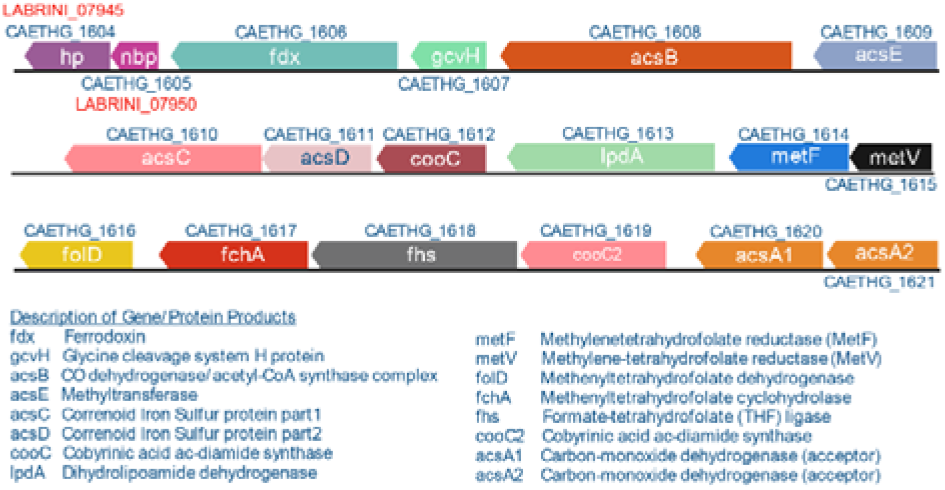
C_1_-fixing gene cluster in Clostridium autoethanogenum. The gene cluster contains 16 genes, including five genes with unconfirmed biochemical functions (CAETHG_1606; 1607; 1612; 1613; 1619). Gene names and annotations based on Valgepea et al., 2022. Figure created using BioRender.com.

Determination of gene functionalities in acetogens through genotype-phenotype studies has been limited mostly due to slow, labour-intensive, and challenging workflows for constructing genetically modified strains (Pavan et al., 2022). However, gene-editing tools for acetogens have rapidly improved recently (Bourgade et al., 2021; Jin et al., 2020; Pavan et al., 2022). We thus aimed to take advantage of these developments to shed light on the functionalities of the *hp* (CAETHG_1604) and the CooT family nickel-binding gene *nbp* (CAETHG_1605), referred to as LABRINI_07945 for *hp* and LABRINI_07950 for *nbp* based on the *C. autoethanogenum* base strain “LAbrini” used in this work (Ingelman et al., 2023).

Here, we used CRISPR/nCas9 in *C. autoethanogenum* for the targeted deletion of two genes adjacent to the C_1_-fixing gene cluster with unclear functions: named *hp* (LABRINI_07945) and *nbp* (LABRINI_07950). Phenotypic characterisation in heterotrophic and autotrophic batch and autotrophic bioreactor continuous cultures revealed significant growth defects and altered by-product profiles for both deletion strains compared to the base strain LAbrini. Additionally, proteomics showed that disruption of either gene affects the expression of proteins of the C_1_-fixing gene cluster and ethanol synthesis pathways. Our findings contribute to a better understanding of genotype-phenotype relationships which is needed to advance the metabolic engineering of acetogen factories for gas fermentation.

## 2. MATERIALS & METHODS

### Bacterial strains, growth media and cultivation conditions

A derivate of *C. autoethanogenum* DSM 10061 strain – DSM X – deposited in the German Collection of Microorganisms and Cell Cultures (DSMZ) named “LAbrini” (Ingelman et al., 2023), was stored as a glycerol stock at −80°C and used as the base strain for genetic engineering. *Escherichia coli* strains NEB Turbo and NEB Express (New England BioLabs) were used for cloning, plasmid assembly, and propagation. For transformant selection, the cultivation of *E. coli* was done in Lysogeny Broth (LB) medium supplemented with ampicillin (50 μg/mL).

For *C. autoethanogenum* cultivation on solid medium, YTF agar (F. Liew et al., 2017) was used with 5 g/L of fructose and antibiotic supplementation if needed (see below). For liquid batch cultures, cells were grown in chemically defined PETC-MES medium (Valgepea et al., 2017) with or without yeast extract (YE; 1.5 g/L) and supplemented with 0.4 g/L of cysteine-HCl·H_2_O as the reducing agent while the carbon source was either 5 g/L of fructose or syngas (50% CO, 20% H_2_, 20% CO_2_, and 10% Ar; AS Eesti AGA). Batch fermentations were performed in 250 mL Schott bottles with 50 mL liquid volume in biological triplicates incubated horizontally at 37 °C with orbital shaking at 120 RPM under strictly anaerobic conditions unless stated otherwise. For autotrophic batch cultures, bottle headspace was pressurised to 140 kPa with syngas. Cultures were sampled during the exponential growth phase for growth characterisation and extracellular metabolome analysis to determine the maximum specific growth rate (μ_max_) and production yields of acetate, ethanol, and 2,3-butanediol. At least five time points were used for μ_max_ calculation resulting in correlation coefficients R^2^ ≥ 0.98 between cultivation time and natural logarithm (ln) of the culture’s optical density (OD_600_). Product yields (mmol/gram of dry cell weight [DCW]) were estimated by linear regression between the respective product concentration (mmol/L) and biomass concentration (gDCW/L) with R^2^ ≥ 0.92.

Continuous chemostat fermentations were carried out as described in detail before (Valgepea et al., 2017a). Shortly, all three *C. autoethanogenum* strains were grown in bioreactors on syngas under strictly anaerobic conditions at 37°C and pH 5 in chemically defined medium (without YE) at a dilution rate of 1 day^-1^. Chemostat continuous cultures were performed in 1.4 L Multifors bioreactors (Infors AG) at a working volume of 750 mL connected to a Hiden HPR-20-QIC mass spectrometer (Hiden Analytical) for online high-resolution off-gas analysis. Steady-state results were collected after OD, gas uptake and production rates had been stable for at least 3–5 working volumes.

### Genetic engineering

#### Genomic and plasmid DNA manipulations

Routine *C. autoethanogenum* genomic DNA extraction for PCR diagnostics was done with PureLink Genomic DNA extraction kit (Invitrogen™, Thermo Fischer Scientific), while plasmid DNA was extracted from *E. coli* using FavorGen FavorPrep™ Plasmid DNA Extraction Mini Kit (Favorgen Biotech Corp). All purified genomic and plasmid DNA fragments were quantified using the NanoDrop™ 1000 Spectrophotometer (Thermo Fischer Scientific). PCR for amplification of DNA fragments for sequencing and cloning, routine screening, and analytical procedures was performed using the Phusion™ High-Fidelity DNA Polymerase (Thermo Fischer Scientific). All primers used in this study (see Supplementary Table 1) were designed using the NetPrimer software (PREMIER Biosoft International) and synthesised by Integrated DNA Technologies. PCR products and DNA fragments were purified from agarose gels with FavorPrep Gel/PCR Purification kit (Favorgen Biotech Corp).

#### Assembly of CRISPR/nCas9 plasmids

A CRISPR/nCas9 gene editing plasmid for *C. autoethanogenum* was constructed based on the pNickClos2.0 plasmid (Addgene plasmid #73228) (Li et al., 2016) with the following modifications to the backbone sequence: origin of transfer and the required traJ coding sequence from pFX01 (Xia et al., 2020) was added to allow potential transfer of the resulting plasmids into recipient cells *via* conjugation. Additionally, one of the two NotI restriction sites was removed to enable convenient cloning of the homology arms (HAs) using NotI and XhoI sites.

To specify the target site for the nCas9 enzyme, single guide RNAs (sgRNAs) were designed using CRISPR RGEN Tools (Park et al., 2015). We used the Basic Local Alignment Search Tool (Altschul et al., 1990) to cross-check that the selected spacer sequences would lead to unique cuts in the genome, thus minimising off-target cleavages. Inverse PCR (iPCR) was employed to clone the unique sgRNA for each target gene into the backbone plasmid to create intermediate plasmids: pGFT021 for LABRINI_07945 (*hp*) and pGFT022 for LABRINI_07950 (*nbp*). The two HAs were obtained by PCR amplification of the 1 kb regions upstream and downstream of each target gene from the *C. autoethanogenum* genome using various primer pairs (Supplementary Table 1). The derived 5’ and 3’ HA fragments were fused using overlap extension PCR as described by (Bryksin and Matsumura, 2013) and moved into the appropriate sgRNA-containing vectors using the same technique or via restriction cloning. See Supplementary File 1 (pGFT036 for *hp*) and 2 (pGFT048 for *nbp*) for plasmid maps. All plasmids constructed in this study are listed in Supplementary Table 2.

#### Plasmid transformation into *C. autoethanogenum* using electroporation

Before electroporation, plasmids were propagated and isolated from the *E. coli* NEB Express strain to lose the Dcm methylation pattern, which has been shown to be targeted by the native type IV restriction system of *C. autoethanogenum* (Woods et al., 2019). Preparation of electrocompetent *C. autoethanogenum* cells and transformation using electroporation was performed as previously described (Leang et al., 2013). Electroporated cells were recovered in 10 mL YTF medium for 18⍰24 h at 37°C with 120 RPM of agitation. Recovery cultures were mixed with 1.5% molten agar YTF supplemented with clarithromycin (4 μg/mL). Transformation plates were incubated inside Oxoid™ AnaeroJars™ (Thermo Fischer Scientific) at 37 °C until colonies became visible (in less than 10 days).

#### Transformant colony screening and plasmid curing

Colonies were first inoculated into liquid YTF medium containing clarithromycin and then plated on selective YTF agar to isolate individual transformant colonies. Next, transformant colonies were randomly chosen for colony PCR screening to verify *C. autoethanogenum* (primers *acs*B-F and -R), plasmid presence (primers oriT-F and -R), and successful gene deletion. Primers *hp*-F, *hp*-R and, *nbp*-F, *nbp*-R were used to confirm deletions of *hp* and *nbp* genes, respectively. PCR products obtained with the gene deletion control primers were purified and verified by Sanger sequencing.

Plasmid-carrying *C. autoethanogenum* colonies with target gene deletions were then subcultured three or more times in 10 mL YTF medium without the antibiotic before plating on non-selective YTF agar to isolate individual colonies. Next, colonies were randomly chosen and patch-plated on both YTF agar without and with clarithromycin (4 μg/mL). Plasmid loss in colonies that failed to grow in the presence of clarithromycin was confirmed by colony PCR using primers oriT-F and oriT-R as above. In total, 10 out of 10 colonies for *hp* gene deletion and 1 out of 30 colonies for *nbp* gene deletion had lost their plasmid. The plasmid-cured colonies were then inoculated into liquid YTF medium for outgrowth and preparation of glycerol stocks, stored as Δ*hp* and Δ*nbp*.

### Analytical Methods

#### Biomass Concentration Analysis

Biomass concentration (gDCW/L) was estimated by measuring culture OD at 600 nm and using the correlation coefficient of 0.23 between OD and DCW using the methodology established before (Peebo et al., 2014).

#### Extracellular Metabolome Analysis

Extracellular metabolome analysis from filtered broth samples was performed as described before (Valgepea et al., 2017b). We note that cells produced 2R,3R-butanediol.

#### Bioreactor Off-Gas Analysis

Bioreactor off-gas analysis for the determination of specific gas uptake (CO and H_2_) and production rates (CO_2_, ethanol) (mmol/gDCW/h) have been described before (Valgepea et al., 2017a). The Faraday Cup detector monitored the intensities of H_2_, CO, ethanol, H_2_S, Ar, and CO_2_ at 2, 14, 31, 34, 40, and 44 amu, respectively.

#### Carbon Balance Analysis

Carbon recoveries and balances were determined as described before (Valgepea et al., 2017a). Briefly, carbon balancing of substrate carbon (CO, cysteine) between growth products (acetate, ethanol, 2,3-BDO, CO_2_, biomass) was calculated as C-mol fraction of each product from total C-mol carbon uptake. Carbon recoveries were calculated as the fraction of summed C-mol products from total C-mol substrates. Ethanol stripping and the total soluble CO_2_ fraction in culture broth were also considered to achieve more accurate carbon balancing.

### Proteome analysis

#### Sample preparation

Proteome analysis was carried out for four biological replicate cultures for each of the three strains: LAbrini, Δ*hp*, and Δ*nbp. C. autoethanogenum* chemostat cultures were sampled for proteomics by immediately pelleting 2 mL of the culture by centrifugation (25,000 × g for 1 min at 4°C) and stored at ⍰80°C until analysis. Cell pellets were suspended with 10 volumes (relative to pellet volume) of chaotrope-based lysis buffer (6 M guanidine-HCl, 100 mM Tris-HCl pH 8.0, 50 mM dithiothreitol), heated at 95°C for 10 min and sonicated with the Bioruptor sonicator (Diagenode) for 15 cycles (30 s ON, 60 s OFF; “High” setting) at 4°C. Next, 0.5 volumes (relative to lysate volume) of Silibeads TypZY-s 0.4⍰0.6 mm (Sigmund Lindner) beads were added to the lysate and bead beating was carried out with the FastPrep24 device (MP Biomedicals) by 2 × 40 s at 6 m/s. Next, samples were centrifuged at 17,000 × g for 5 min, and the supernatant was separated from the beads and transferred to a new tube. A small aliquot of the lysate was then precipitated with trichloroacetic acid-deoxycholate (TCA-DOC) precipitation and the protein concentration was determined with the Micro BCA assay (Thermo Fisher Scientific). Next, 20 μg of lysate protein was alkylated with 100 mM chloroacetamide by incubating for 1 h in the dark at room temperature. Samples were then processed, and proteins were digested to peptides by the SP3 protocol as described elsewhere (Hughes et al., 2018).

#### LC-MS/MS analysis

Sample order for LC-MS/MS analysis was randomised to avoid batch effects and blanks were flanked to minimise carry-over. 1 μg of peptides for each sample was injected to an Ultimate 3500 RSLCnano system (Dionex) using a 0.3 × 5 mm trap-column (5 μm C18 particles, Dionex) and an in-house packed (3 μm C18 particles, Dr Maisch) analytical 50 cm x 75 μm emitter-column (New Objective). Columns were operated at 45°C. Peptides were eluted at 300 nL/min with an 8-45% B 90 min gradient (buffer B: 80% acetonitrile + 0.1% formic acid, buffer A: 0.1% formic acid) to a Q Exactive HF (Thermo Fisher Scientific) mass spectrometer (MS) using a nano-electrospray source (spray voltage of 2.5 kV in positive mode). The MS was operated using a data-independent acquisition (DIA) approach (Gillet et al., 2012) with variable isolation windows over a range of 400⍰1,100 m/z. Briefly, one full range 400⍰1,100 m/z MS1 scan was collected at a resolution setting of 60,000 (max ion injection time of 60 ms, max of 3e6 ions), followed by 25 overlappings (overlap of 1 m/z) DIA isolation windows as follows (start□end m/z, isolation width): 399⍰472, 25; 47⍰510, 20; 509⍰608, 15; 607⍰703, 20; 702⍰790, 30; 789⍰868, 40; 867⍰966, 50; and 965⍰1101, 136. Each DIA scan was collected at a resolution setting of 30,000 (max ion injection time of 41 ms, max of 3e6 ions), the default charge state was set to +3 and normalised collision energy to 27. The overall method cycle time was approximately 2.3 s.

#### DIA MS data analysis and differential protein expression analysis

DIA MS data was analysed using version 1.8 of the software suite DIA-NN (Demichev et al., 2019) with default settings. The spectral library was generated *in silico* using smart profiling from the protein sequence database of *C. autoethanogenum* LAbrini (NCBI Genbank CP110420) (Ingelman et al., 2023) with a precursor of 1% false discovery rate (FDR) filter. The following settings were used for analysis: full trypsin specificity with 1 missed cleavage allowed for peptides with a length of 7–30 AAs; fixed modifications of cysteine carbamidomethylation and methionine N-terminal excision; precursor charge range 1–4 and m/z range 400–1,100; fragment m/z range 200–1,800; double-pass mode for neural network classifier; Robust LC (high precision) for quantification strategy; and RT-dependent cross-run normalization. Label-free protein quantification and normalisation were performed within the DIA-NN workflow using the MaxLFQ method (Cox et al., 2014). We confidently quantitated 28,752 peptides and 2,127 proteins across all samples, and 27,358 peptides and 2,107 proteins on average within each sample after the removal of shared peptides from the analysis.

Protein expression fold-changes (FC) with q-values between Δ*hp* and LAbrini, and Δ*nbp* and LAbrini were determined using the software Perseus (Tyanova et al., 2016) with Student’s T-test. Only proteins with at least two peptides in all 12 samples (1,835) analysed were used to ensure higher quantification accuracy. Proteins were considered to be differentially expressed by an FC > 1.5 and a q-value < 0.05 after FDR correction (Benjamini and Hochberg, 1995). Differentially expressed proteins are presented in Supplementary Table 3. Proteomics data have been deposited to the ProteomeXchange Consortium (http://proteomecentral.proteomexchange.org) via the PRIDE partner repository (Perez-Riverol et al., 2022) with the data set identifier PXDYYY.

## 3. RESULTS AND DISCUSSION

### Single-gene deletion of *hp* and *nbp* in *C. autoethanogenum* using CRISPR/nCas9 knock-out plasmids

To study the role of two genes adjacent to the C_1_-fixing gene cluster (Figure 1) with unclear functions, LABRINI_07945 (*hp*) and LABRINI_07950 (*nbp*), each gene was targeted for deletion using CRISPR/nCas9. Plasmids pGFT036 and pGFT048 carrying required components (Figure 2A) for *hp* and *nbp* deletion, respectively, were transferred into *C. autoethanogenum* by electroporation that yields transformation efficiencies up to 200 CFU/μg DNA in our laboratory. We obtained 40 and 2 colonies for *hp* and *nbp* deletion, respectively, and colony PCR screening of 6 colonies for *hp* and 2 for *nbp* deletion confirmed *C. autoethanogenum* genotype and the presence of our relatively large plasmid (∼12.5 kb) in all colonies. The gene deletion control PCR showed that all the *hp* colonies screened had a mixed base strain/knock-out (KO) genotype evidenced by the presence of both the 2444-bp fragment of the base strain allele and the predicted 2103-bp fragment of the KO allele (Figure 2B). Six rounds of liquid selective sub-culturing of the mixed colonies was needed to obtain 1 colony (from 20 screened) with a clean *hp* KO genotype (Figure 2C). In contrast, the 2 *nbp* colonies (*nbp*-C1 and *nbp*-C2) obtained showed a 1444-bp fragment of the desired KO allele, i.e. a clean KO genotype (Figure 2B). Sanger sequencing confirmed both *hp* and *nbp* gene deletions in selected colonies and the presence of a PstI site, proving that the provided plasmid-based DNA repair template, in the form of the fused 1 kb HA interspaced with the PstI site, was used during homologous recombination to repair the nick from nCas9 (Figure 2C). Successful deletion of both genes was also confirmed by proteome analysis (see below). Finally, the obtained Δ*hp* and Δ*nbp* cells were cured of the plasmid before proceeding with phenotypic characterisation.

**Figure 2.**
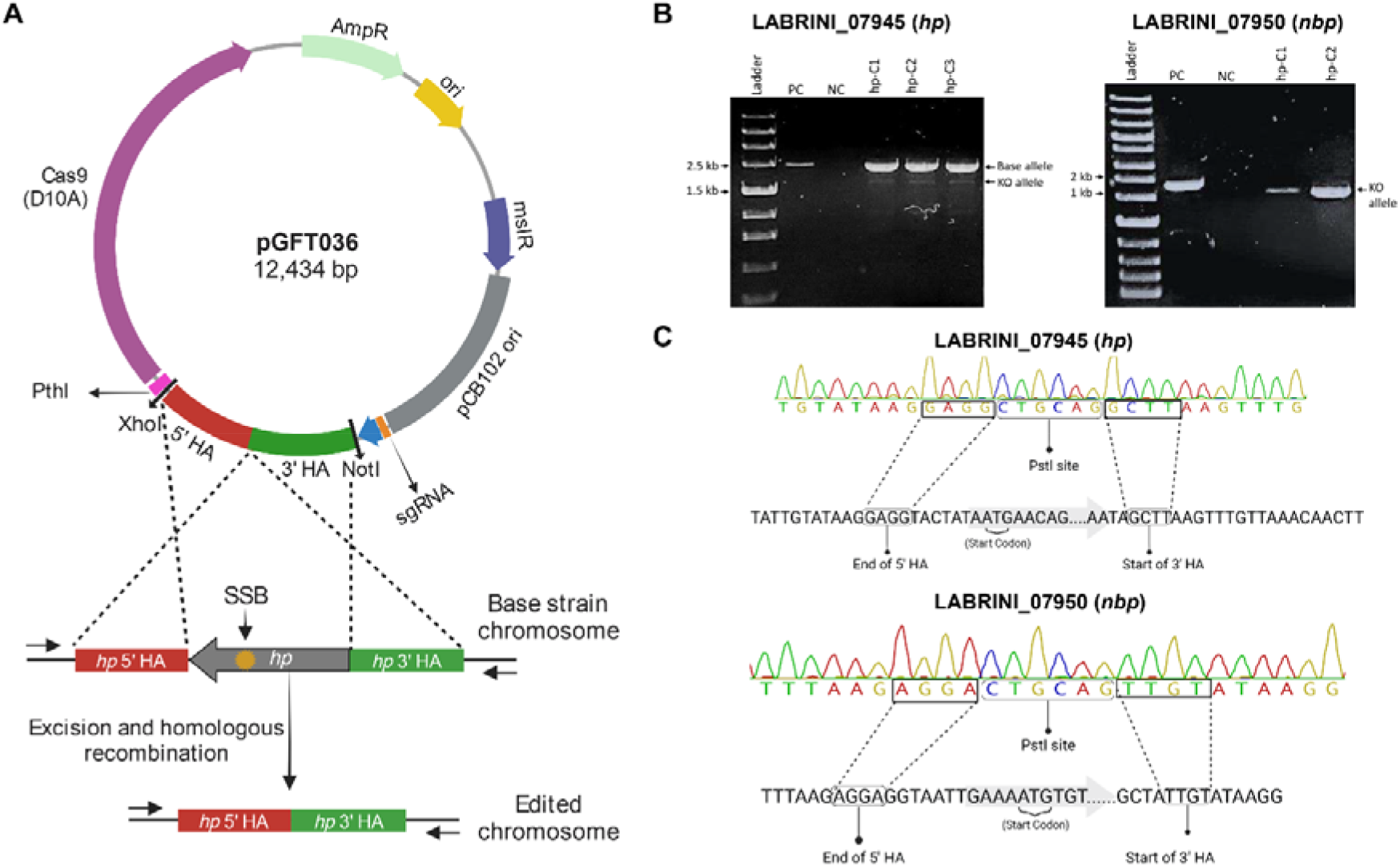
Successful deletion of target genes using CRISPR/nCas9 KO plasmids in *C. autoethanogenum*. A. Representative plasmid map and an overview of CRISPR/nCas9 mediated gene deletion mechanism. The yellow star indicates the genomic recognition site of the Cas9-sgRNA complex. B. PCR Screening of transformant colonies using control primers to amplify gene regions. C. Confirmation of *hp* (LABRINI_07945) and *nbp* (LABRINI_07950) deletion via Sanger Sequencing. PC, positive control (non-transformed *C. autoethanogenum*); NC, negative control (no template); LAbrini, *C. autoethanogenum* base strain allele; KO, knock-out/deletion allele; SSB, single-strand break. Figure partly created using BioRender.com.

### Deletion of *hp* and *nbp* genes confer growth defects in rich-medium batch cultures

Next, we grew Δ*hp* and Δ*nbp* strains in batch cultures on PETC-MES medium with YE (i.e. rich medium) in heterotrophic (fructose) and autotrophic (syngas: CO+H_2_+CO_2_) conditions to investigate potential phenotypic effects of *hp* and *nbp* deletions. Interestingly, the maximum specific growth rate (μ_max_) of both mutant strains was significantly impaired compared to the *C. autoethanogenum* base strain “LAbrini” on both fructose and syngas (Figure 3A). The μ_max_ for the Δ*hp* strain was 0.08 ± 0.001 (average ± standard deviation) and 0.05 ± 0.001 on fructose and syngas, respectively, while for the Δ*nbp* strain, the μ_max_ was 0.07 ± 0.002 on fructose and 0.06 ± 0.001 on syngas (Figure 3A). The LAbrini base strain showed μ_max_ values of 0.10 ± 0.001 on fructose and 0.08 ± 0.001 on syngas. Thus, μ_max_ for deletion strains were substantially lower compared to base strain: for Δ*hp*, 21% and 37% lower on fructose and syngas and for Δ*nbp*, 34% and 26% lower on fructose and syngas, respectively (Figure 3A). Interestingly, Δ*hp* had a more pronounced growth defect on syngas, while Δ*nbp* was affected more on fructose. The μ_max_ values for both growth conditions were also statistically different between deletion strains. In addition to the lower μ_max_, Δ*hp* strain exhibited a longer lag phase and slightly lower peak OD on syngas compared to the base strain LAbrini (Figure 3B). The growth profile and peak OD of the *Δnbp* strain, however, were similar to LAbrini despite the lower μ_max_ values (Figure 3C). Altogether, our results demonstrate that the deletion of either the *hp* or the *nbp* gene confers significant growth defects for hetero- and autotrophic rich medium batch cultures, suggesting a notable role for both genes in the physiology of *C. autoethanogenum*.

**Figure 3.**
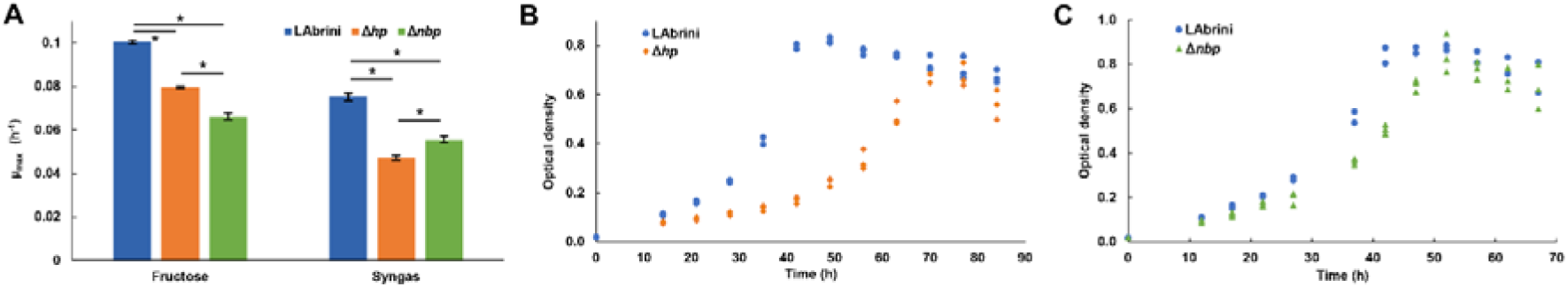
Batch growth of Δ*hp*, Δ*nbp*, and LAbrini strains on PETC-MES with YE. A. Maximum specific growth rate (μ_max_) in heterotrophic (fructose) and autotrophic (syngas) growth conditions. Data are average ± standard deviation between three biological replicates. Asterisk denotes values statistically different according to t-test (p-value ≤ 0.01). B and C. Autotrophic growth profiles on syngas. LAbrini, *C. autoethanogenum* base strain; YE, yeast extract.

### Deletion of *hp* and *nbp* show no effects on growth in minimal medium autotrophic batch cultures

We also investigated the growth of all three strains in PETC-MES medium without YE to assess the effects of *hp* and *nbp* deletions on autotrophic growth on syngas in minimal medium batch cultures. Intriguingly, there was no difference in the μ_max_ values between the three strains: 0.08 ± 0.004, 0.08 ± 0.005, and 0.08 ± 0.003 for the Δ*hp*, Δ*nbp*, and LAbrini strains, respectively (Figure 4A). Although one would expect slower growth in minimal medium, Δ*hp* and Δ*nbp* strains grew faster in minimal medium compared to rich medium, whereas LAbrini showed similarly high μ_max_ values for autotrophic growth of acetogens on both media. Like the μ_max_ data, the three strains did not show significant variations in peak OD and growth curves (i.e. lag phase and entry into stationary phase) (Figure 4B). Thus, both genes seem to play a role in *C. autoethanogenum* for utilisation of complex nutrients due to significant growth defects in rich medium with no phenotypic effects in minimal medium for Δ*hp* and Δ*nbp* strains.

**Figure 4.**
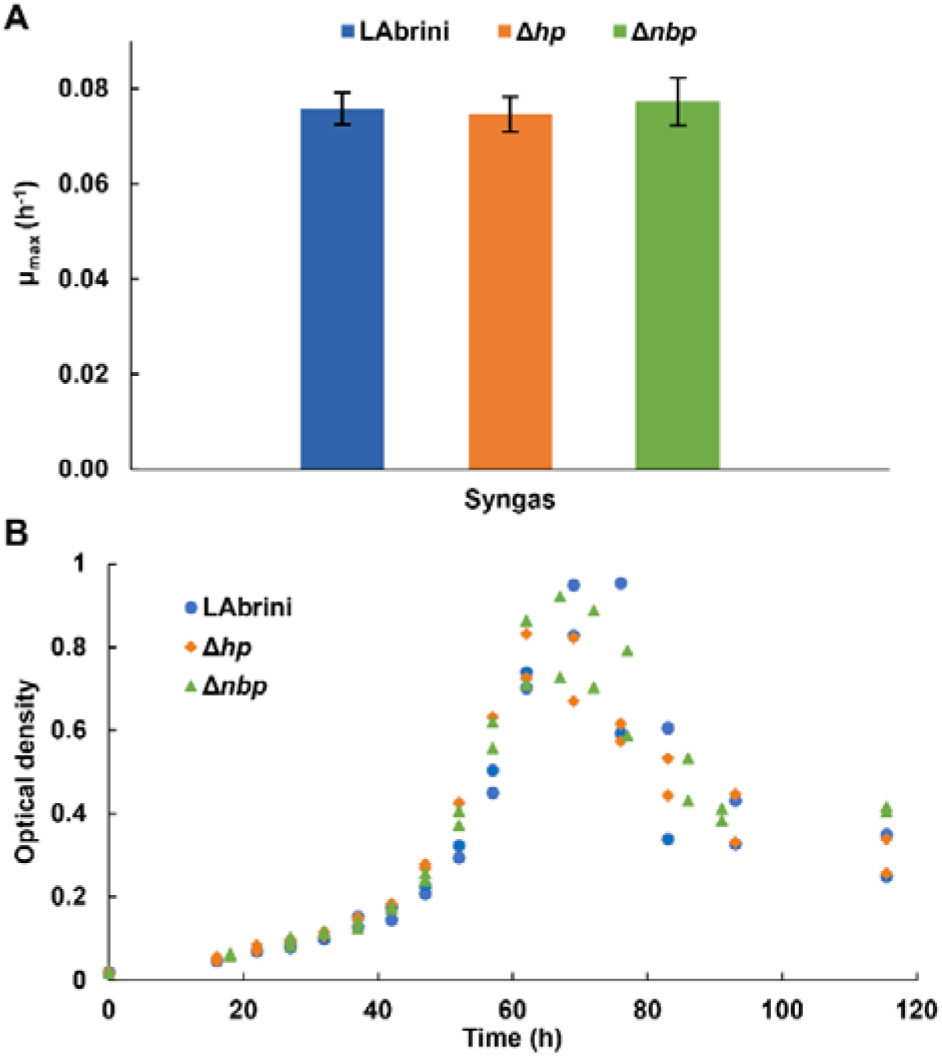
Autotrophic batch growth of Δ*hp*, Δ*nbp*, and LAbrini strains in PETC-MES without YE. A. Maximum specific growth rates (μ_max_) on syngas. Data are average ± standard deviation between two biological replicates. B. Growth profiles on syngas. LAbrini, *C. autoethanogenum* base strain; YE, yeast extract.

### Deletion of *hp* and *nbp* alter growth by-product yields in autotrophic batch cultures

We determined the production yields (mmol/gDCW) of *C. autoethanogenum* growth by-products – ethanol, acetate, and 2,3-butanediol (2,3-BDO) – during the above-described autotrophic batch cultures to check if these are affected by the deletion of either the *hp* or the *nbp* gene. Remarkably, the Δ*hp* strain showed a >5-fold higher acetate yield (550 ± 45 mmol/gDCW) relative to LAbrini (92 ± 7 mmol/gDCW) during growth in rich medium (Figure 5A). The production of 2,3-BDO was also slightly higher for Δ*hp* compared to LAbrini (10 ± 1 vs 7 ± 1 mmol/gDCW). While the ethanol yield was not different for Δ*hp* when compared to LAbrini (96 ± 5 vs 102 ± 13 mmol/gDCW), ethanol production did not coincide with acetate production in the Δ*hp* strain (data not shown). Namely, Δ*hp* was producing acetate in the early exponential growth phase and started producing ethanol only in the late exponential phase.

**Figure 5.**
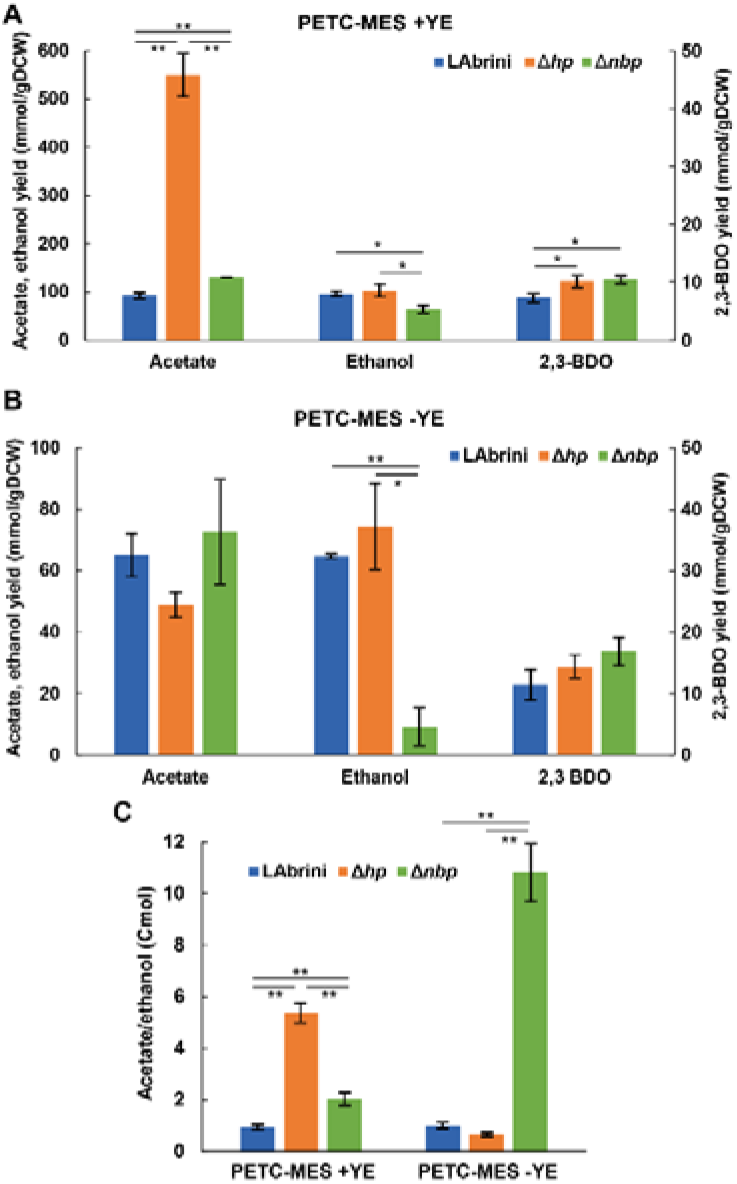
Growth by-product yields and acetate-to-ethanol ratios of Δ*hp*, Δ*nbp*, and LAbrini strains in autotrophic batch cultures with and without YE. A and B. Growth by-product yields. C. Acetate/ethanol (Cmol). Data are average ± standard deviation between three biological replicates on PETC-MES +YE or two biological replicates on PETC-MES -YE. Asterisk denotes values statistically different according to t-test: *, p-value < 0.05; **, p-value < 0.01. LAbrini, *C. autoethanogenum* base strain; YE, yeast extract; gDCW, gram of dry cell weight; 2,3-BDO, 2,3-butanediol.

Also, the Δ*nbp* strain showed significantly different product yields compared to LAbrini during growth in rich medium: 41% higher acetate (130 ± 0.2 vs 92 ± 7 mmol/gDCW), 44% higher 2,3-BDO (10 ± 1 vs 7 ± 1 mmol/gDCW), and 33% lower ethanol compared to LAbrini (65 ± 8 vs 96 ± 6 to mmol/gDCW) (Figure 5A). In addition, both acetate and ethanol production yields were different between the two deletion strains. The substantially higher acetate production yield of the Δ*hp* strain might indicate that deletion of the *hp* gene improves the uptake of YE components and their catabolism to acetate. However, this did not lead to faster growth, thus, acetate production was potentially not linked to ATP production.

We observed no difference in the product yields between the Δ*hp* strain and LAbrini on minimal medium (Figure 5B), which is not surprising as growth was also comparable (Figure 4). However, despite similar growth for Δ*nbp* and LAbrini strains on minimal medium (Figure 4), the Δ*nbp* strain showed an 86% lower ethanol yield than LAbrini (9 ± 6 vs 65 ± 1 to mmol/gDCW) while acetate and 2,3-BDO yields were similar (Figure 5B). Only the ethanol production yield was different between the deletion strains on minimal medium. Interestingly, while LAbrini maintained the acetate-to-ethanol ratio at ∼1 on both media, the ratio was higher for the Δ*hp* strain (∼5 vs ∼1) and lower for the Δ*nbp* strain (∼2 vs ∼11) on rich medium compared to minimal medium during autotrophic growth (Figure 5C). These ratios are in the range of those previously seen in autotrophic batch and continuous bioreactor cultures of *C. autoethanogenum* (Ingelman et al., 2023; Valgepea et al., 2017a, 2018). Our results imply that both *hp* and *nbp* genes play a notable role in carbon distribution between the two major by-products – acetate and ethanol – during autotrophic batch growth of *C. autoethanogenum*.

### Deletion of *hp* and *nbp* affect growth characteristics of steady-state chemostat cultures

We also compared the performance of the mutant strains to the LAbrini base strain in steady-state bioreactor continuous cultures as these yield high-quality physiological data (Adamberg et al., 2015; Valgepea et al., 2018) and are also the industrially relevant fermentation mode for acetogens (Köpke and Simpson, 2020). Cells were grown in minimal medium in syngas chemostats at a dilution rate of 1 day^-1^ and steady-state data at biomass concentrations ∼1.5 gDCW/L were collected for by-product production (ethanol, acetate, 2,3-BDO) and gas uptake and production (CO, H_2_, CO_2_) analysis.

While acetate production in minimal medium batch cultures was similar across all three strains, both Δ*hp* and Δ*nbp* strains showed lower specific acetate production rates (q_ace_; mmol/gDCW/day) in chemostats (Figure 6A). Specific ethanol production rates (q_EtOH_) were also lower for both deletion strains compared to LAbrini (Figure 6A), whereas ethanol production differed only for the Δ*nbp* strain in batch cultures (Figure 5). Production of 2,3-BDO (q_2,3-BDO_) was similar across the three strains in chemostats (Figure 6A). These steady-state data are consistent with above-described batch data by showing that both *hp* and *nbp* genes affect carbon distribution during autotrophic growth of the model-acetogen *C. autoethanogenum*. Still, differences in bioreactor chemostat and bottle batch data demonstrate the importance of testing strains in the industrially relevant fermentation mode for acetogens (Köpke and Simpson, 2020).

**Figure 6.**
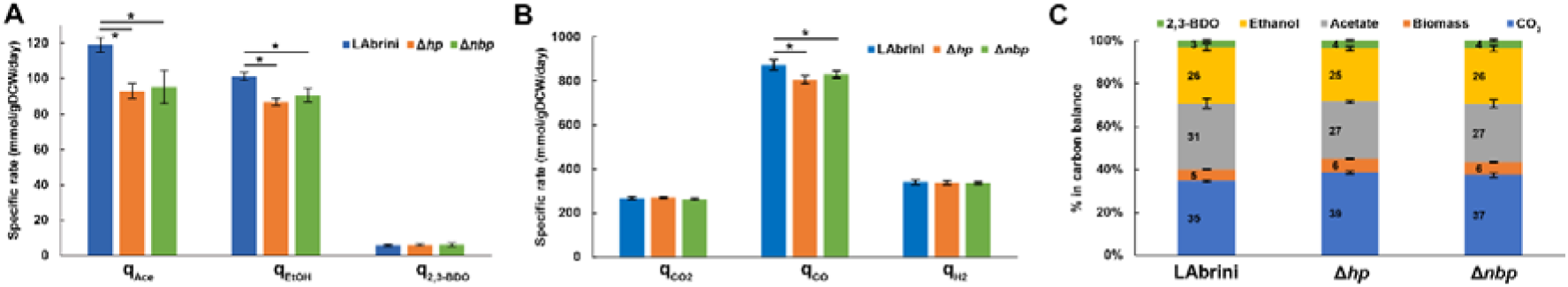
Steady-state growth characteristics of Δ*hp*, Δ*nbp*, and LAbrini strains in syngas-grown chemostats. A. Specific by-product production rates (mmol/gDCW/day). B. Specific gas uptake (CO, H_2_) and production (CO_2_) rates. Asterisk denotes values statistically different according to t-test (p-value < 0.05). C. Carbon balances. Carbon recoveries were normalised to 100% to have a fair comparison of carbon distributions between different strains. Data are average ± standard deviation between four biological replicates. LAbrini, *C. autoethanogenum* base strain; EtOH, ethanol; 2,3-BDO, 2,3-butanediol; Ac, acetate; gDCW, gram of dry cell weight; *q*, specific rate.

Bioreactor off-gas analysis revealed that the specific CO uptake rates (q_CO_; mmol/gDCW/day) were slightly lower for both deletion strains compared to LAbrini (873 ± 19): by 8% for the Δ*hp* strain (805 ± 19) and by 5% for the Δ*nbp* strain (830 ± 15) (Figure 6B). At the same time, we detected no differences in the specific H_2_ uptake rates (q_H2_; mmol/gDCW/day) and CO_2_ production rates (q_CO2_; mmol/gDCW/day) among the three strains (Figure 6B). The slightly higher q_CO2_/q_CO_ and q_H2_/q_CO_ for both deletion strains indicates that cells needed to increase the generation of reducing power in the form of reduced ferredoxin as a response to gene deletion since both CO and H_2_ oxidation supply reduced ferredoxin (Bertsch and Müller, 2015; Pavan et al., 2022). Quantification of carbon flows from substrates to products showed that both Δ*hp* and Δ*nbp* strains diverted more carbon to biomass and CO_2_ with a concomitant lower carbon flux to ethanol and acetate (Figure 6C). The slightly lower acetate-to-ethanol ratios for Δ*hp* (1.2) and Δ*nbp* (1.1) compared to LAbrini (1.3) are consistent with higher q_CO2_/q_CO_ for deletion strains since ethanol production generally results in a greater loss of carbon from CO as CO_2_ (Bertsch and Müller, 2015).

### Proteome analysis reveals intriguing effects of deletion of *hp* and *nbp* genes

We used proteomics to determine the effects of *hp* and *nbp* gene deletions on protein expression in the above-described chemostat cultures of *C. autoethanogenum*. Our analysis quantified 93 differentially expressed proteins (fold-change > 1.5 and a q-value < 0.05) between *hp* and LAbrini, and 77 between *nbp* and LAbrini (Figure 7; Supplementary Table 3). Proteomics also confirmed the deletion of each gene in respective strains as no peptides for either protein in corresponding strains were detected. Intriguingly, deletion of either gene increased the expression of the methyltransferase (AcsE; LABRINI_07970) and a component of the correnoid iron sulphur protein (AcsD; 07980) (Table 1) that are linked to a critical activity of the WLP: synthesis of acetyl-CoA. AcsE and AcsD supply the methyl group to the carbon monoxide dehydrogenase (AcsA; LABRINI_08025) and acetyl-CoA synthase (AcsB; 07965) CODH/ACS enzyme complex (Lemaire and Wagner, 2021), that is essential for autotrophic growth of *C. autoethanogenum* (Liew et al., 2016). At the same time, the expression of AcsB was repressed in both deletion strains (Table 1). These are all among the top 10 most abundant proteins for autotrophic growth of *C. autoethanogenum*, except AcsA (Valgepea et al., 2022). These results show that the two genes adjacent to the C_1_-fixing cluster studied here – *hp* and *nbp* – affect the expression of key proteins for C_1_-fixation. Notably, the effect of *hp* and *nbp* on C_1_-fixation extends beyond the WLP as the expression of the most abundant CODH in *C. autoethanogenum* (Valgepea et al., 2022) – CooS1; 15015 – was repressed in both mutant strains (Table 1). Further work is needed to understand the interplay between the *hp, nbp*, and the affected enzymes with a central role in the autotrophic growth of *C. autoethanogenum*.

**Table 1.** Key protein expression changes in Δ*hp* and Δ*nbp* strains compared to LAbrini.

| Protein ID | Description of protein product | $\Delta hp$ vs. LABrini | | | $\Delta nbp$ vs. LABrini | | |
| --- | --- | --- | --- | --- | --- | --- | --- |
|  |  | Log<sub>2</sub> FC | FC | q-value | Log<sub>2</sub> FC | FC | q-value |
| *C<sub>1</sub>-fixation* |  |  |  |  |  |  |  |
| LABRINI_07970 | Methyltransferase for CODH | 0.61 | 1.5 | 0.01 | 0.63 | 1.6 | 0.03 |
| LABRINI_07980 | Correnoid iron sulphur protein part2 | 0.58 | 1.5 | 0.01 | 0.63 | 1.5 | 0.02 |
| LABRINI_07965 | Acetyl-CoA synthase (ACS) | -0.60 | -1.5 | < 0.01 | -0.88 | -1.8 | < 0.01 |
| LABRINI_15015 | Carbon-monoxide dehydrogenase CooS1 | -0.55 | -1.5 | 0.02 | -0.83 | -1.8 | 0.01 |
| *Ethanol synthesis* |  |  |  |  |  |  |  |
| LABRINI_09125 | Iron-containing alcohol dehydrogenase | 1.71 | 3.3 | 0.01 | 1.75 | 3.4 | 0.01 |
| LABRINI_19700 | Iron-containing alcohol dehydrogenase | 0.82 | 1.8 | 0.01 | 0.93 | 1.9 | 0.01 |
| LABRINI_18005 | Iron-containing alcohol dehydrogenase | 0.61 | 1.5 | 0.01 | 0.82 | 1.8 | 0.01 |
| LABRINI_00495 | Aldehyde ferredoxin oxidoreductase | 1.10 | 2.1 | 0.03 | 1.47 | 2.8 | 0.03 |
| LABRINI_00445 | Aldehyde ferredoxin oxidoreductase | 0.61 | 1.5 | 0.02 | 0.02 | 1.6 | 0.02 |
| LABRINI_18700 | Bifunctional AdhE1 | 0.77 | 1.7 | 0.01 | 2.07 | 4.2 | < 0.01 |

ID, identifier; FC, fold-change. Positive FC means up-regulation, negative down-regulation. See Supplementary Table 3 for all differentially expressed proteins.

**Figure 7.**
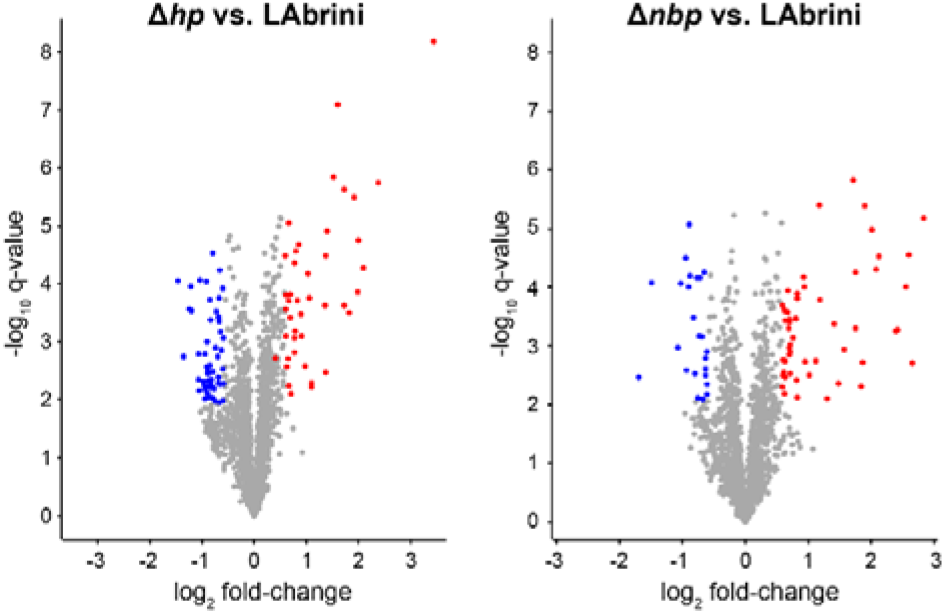
Volcano plots showing differentially expressed proteins in Δ*hp* and Δ*nbp* strains compared to LAbrini. Differentially expressed proteins are with fold-change > 1.5 and a q-value < 0.05. Blue markers, down-regulated proteins; red markers, up-regulated proteins; grey, proteins not expressed differentially. See Supplementary Table 3 for data.

Deletion of our target genes also affected the expression of proteins involved in the synthesis of the most attractive native product of *C. autoethanogenum*: ethanol. In contrast to the lowered ethanol production in the deletion strains, proteome analysis revealed that expression of alcohol dehydrogenases (19700 and 18005) was elevated in both mutant strains, including the most abundant alcohol dehydrogenase in *C. autoethanogenum* (Valgepea et al., 2022): Adh4, 09125 (Table 1). Similarly, increased expression of both aldehyde ferredoxin oxidoreductases (AOR1, 00445 and AOR2, 00495), that couple ethanol synthesis to ATP production and the bifunctional acetaldehyde-CoA/alcohol dehydrogenase AdhE1 (18700) were measured in deletion strains. Such seeming discrepancies between either the transcripts or proteins linked to ethanol synthesis and the measured ethanol production have been detected in *C. autoethanogenum* before (Valgepea et al., 2017a, 2018). Thus, more systemic and in-depth investigations are required to decipher the genotype-phenotype relationships for ethanol synthesis in acetogens.

Among other differentially expressed proteins, it was unexpected to see repression of 13 ribosomal proteins in the Δ*hp* strain (1.7- to 2.3-fold, q < 0.05; Supplementary Table 3). In addition, several oxidoreductases (05235, 14800, and 18000; ∼2-fold, q < 0.03) and sigma-54-related transcriptional regulators (14025 and 00450; ∼3.2- and 1.6-fold, q < 0.05) were up-regulated in the deletion strains. Also noteworthy is the ∼1.8-fold (q < 0.01) up-regulation of a PocR ligand-binding domain-containing protein (16380) in both Δ*hp* and Δ*nbp* strains, as we recently identified a missense mutation in the PocR gene of a CO-evolved *C. autoethanogenum* strain obtained via adaptive laboratory evolution (Ingelman et al., 2023). Moreover, PocR has an alternative annotation in *C. autoethanogenum* as a histidine kinase (Humphreys et al., 2015) that regulates sporulation and metabolism in Clostridia (Xin et al., 2020).

## 4. CONCLUSION

In summary, our study shows that CRISPR/nCas9 can effectively be used for targeted gene deletion in the industrially relevant acetogen *C. autoethanogenum* and provides valuable insights to accelerate the rational genetic engineering of acetogen cell factories. Additionally, the characterisation of the constructed deletion strains offers valuable clues to the *in vivo* functionalities of the *hp* and *nbp* genes that seem to influence both growth and product patterns of *C. autoethanogenum* during autotrophic growth. Furthermore, our proteomics results shed light on enzymes that potentially play a key role in ethanol production. Future work will need to confirm the biochemical function of closely linked genes in the WLP and identify possible targets for improving carbon fixation efficiencies.

## ACKNOWLEDGEMENTS

We thank Dr Bastian Molitor and Dr Peng-Fei Xia for providing us the pFX01 plasmid.

## AUTHOR CONTRIBUTIONS

Conceptualization: UJN, KR, and KV; Methodology: UJN, KR, KMS, AH, MK, and KV; Formal analysis: UJN, KR, LAL, PRP, and KV; Investigation: UJN, KR, LAL, PRP, and KV; Resources: KV; Writing – Original Draft: UJN, KR, and KV; Writing – Review & Editing: UJN, KR, LAL, PRP, KMS, AH, MK, and KV; Supervision: KV; Project Administration: KV; Funding Acquisition: KV.

## DATA AVAILABILITY STATEMENT

Proteomics data have been deposited to the ProteomeXchange Consortium (http://proteomecentral.proteomexchange.org) via the PRIDE partner repository (Perez-Riverol et al., 2022) with the dataset identifier PXDYYY.

